# Direct analysis of ribosome targeting illuminates thousand-fold regulation of translation initiation

**DOI:** 10.1101/2020.04.28.066068

**Authors:** Rachel O. Niederer, Maria F. Rojas-Duran, Boris Zinshteyn, Wendy V. Gilbert

## Abstract

Translational control shapes the proteome in normal and pathophysiological conditions. Current high-throughput approaches reveal large differences in mRNA-specific translation activity but cannot identify the causative mRNA features. We developed direct analysis of ribosome targeting (DART) and used it to dissect regulatory elements within 5′ untranslated regions that confer thousand-fold differences in ribosome recruitment in biochemically accessible cell lysates. Using DART, we identified novel translational enhancers and silencers, determined a functional role for most alternative 5′ UTR isoforms expressed in yeast, and revealed a general mode of increased translation via direct binding to a core translation factor. DART enables systematic assessment of the translational regulatory potential of 5′ UTR variants, whether native or disease-associated, and will facilitate engineering of mRNAs for optimized protein production in various systems.

**Highlights:**

- DART illuminates thousand-fold differences in 5′ UTR-specific translation activity
- SNPs and alternative 5′ UTR isoforms affect ribosome recruitment significantly
- Inhibitory effects of RNA structures are highly dependent on 5′ UTR context
- 5′ UTR motifs bind initiation factors directly, broadly stimulating translation

## Introduction

Translation initiation is an important step in eukaryotic gene expression whose dysregulation is linked to heritable human diseases and cancer. Systematic characterization has shown that mRNA-specific translational activity varies by orders of magnitude under normal growth conditions and is extensively regulated in response to a wide range of physiological signals^1–6^. Despite growing interest in the mRNA features responsible for this widespread translational control, current high-throughput methods, which rely on quantification of the average number of translating ribosomes per transcript, have revealed only partial correlations that leave many differences in translation activity unexplained^7,8,9^. In contrast, in-depth genetic and biochemical analysis has revealed detailed regulatory mechanisms for certain mRNAs^10,11^.

Genome-wide association studies (GWAS) show diverse disease phenotypes associated with non-coding single-nucleotide polymorphisms (SNPs) found within 5′-UTRs. However, current efforts to identify causal variants, which are potential targets for new therapies, are largely restricted to non-synonymous changes within protein coding sequences. A richer understanding of mRNA sequence-specific translation mechanisms is needed to predict which 5′-UTR variants are likely to be damaging by dysregulating protein levels. We developed DART technology as a bridge between high-throughput, but mechanistically difficult to parse, in vivo approaches and mechanistically precise, but low-throughput, in vitro translation reconstitution assays. In DART, thousands of synthetic mRNAs initiate translation in vitro. Differential abundance analysis of ribosome-bound and input mRNA reveals differences in translation activity. By massively parallel testing of defined sequences, both endogenous and mutated, our approach moves beyond correlation analysis to enable rapid, hypothesis-driven dissection of the causative role of specific 5′-UTR elements to determine their mechanisms of action.

Here, we demonstrate the broad power of DART to illuminate the mechanisms underlying translational control. We tested 4354 alternative mRNA isoforms from 2064 yeast genes revealing widespread translational control by alternative 5′-UTRs. We leveraged these alternative 5′-UTR isoform comparisons to discover hundreds of previously uncharacterized translational enhancers and silencers, short sequence elements that are sufficient to promote or repress ribosome recruitment. We also exploited the throughput of the DART approach to systematically interrogate the effects of RNA secondary structure and start codon context on translation initiation. Our results demonstrate highly context-dependent control by inhibitory 5′-UTR structures. These unanticipated sequence context effects likely explain the weak correlation between predicted RNA folding and translation activity observed previously, as well as the idiosyncratic translational consequences of inactivating mRNA helicases. Finally, we used DART to directly test the function of initiation factor binding motifs within 5′-UTRs. Our results establish a broad stimulatory role for eIF4G1 binding motifs in 5′-UTRs and illustrate the potential for DART to illuminate the function of putative RNA regulatory elements identified by other high-throughput approaches. Together, our results reveal thousands of previously unidentified functional elements within 5′-UTRs that substantially affect translation. This study establishes DART as a powerful new high-throughput method that can be broadly applied to both discover and interrogate regulatory features within 5′-UTRs.

## Results

### Development of DART for quantitative comparison of translation initiation

5′-UTRs directly contact the translation initiation machinery and can strongly influence translation activity (**Fig. 1a**), but the features that distinguish efficiently translated mRNAs are largely unknown. We developed direct analysis of ribosome targeting (DART) as a high-throughput method to enable rapid determination of the role of individual regulatory elements within 5′-UTRs. We designed a library of DNA oligos, based on deep sequencing of yeast mRNA isoforms^12^, that included 12,000 sequences corresponding to 4,252 genes. This library covers 68% of genes expressed during exponential growth and includes alternative mRNA isoforms for many genes. A common (5′) T7 promoter and a (3′) priming site for reverse transcription flanked each unique mRNA sequence, which included the 5′-UTR plus a short region of coding sequence to preserve the endogenous start codon context. Any upstream AUGs within 5′-UTR sequences were mutated so the first AUG encountered by a scanning pre-initiation complex moving 5′ to 3′ would be the annotated AUGi. Each oligo also included a unique 10nt barcode to distinguish closely related sequences (**Fig. 1b**).

**Fig. 1.**
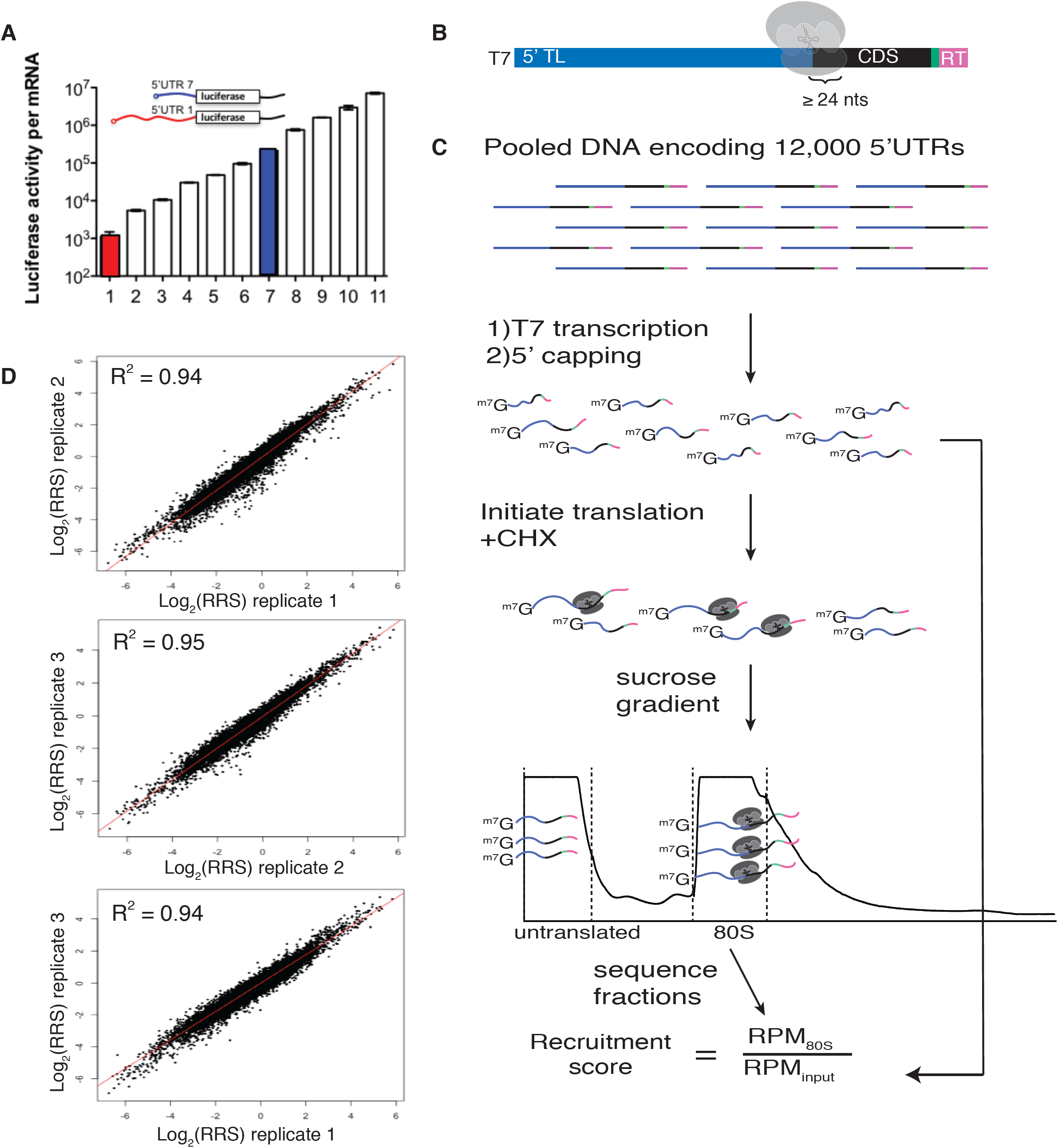
DART quantifies 5′-UTR-mediated translational control over a thousand-fold range. **(A)** 5′-UTR isoforms confer over a thousand-fold range in protein production. 5′-UTRs added to a common luciferase gene result in large differences in translational output. Red and blue bars represent 5′-UTR isoforms from a single gene. **(B)** Schematic of DNA pool design. All sequences begin with the T7 promoter followed by ≤122nts of 5′-UTR sequence, then a minimum of 24nts of coding sequence, followed by a 10nt identifier barcode, and finally an RT handle for library preparation. **(C)** DART-seq overview. **(D)** DART-seq reproducibly measures ribosome recruitment over a thousand-fold range. Comparison of 3 DART-seq replicates.

Next, we used these designed pools to generate dsDNA templates for in vitro transcription followed by enzymatic mRNA capping to produce the pooled RNA substrates for in vitro translation initiation. RNAs were enzymatically biotinylated at their 3′ ends to facilitate quantitative recovery. RNA pools were incubated in yeast translation extracts^13–15^ for 30 min, a time point that maximizes quantitative differences in ribosome recruitment between 5′-UTRs while yielding enough initiated mRNA for robust library preparation. Cycloheximide was included in the translation reaction to freeze recruited ribosomes at initiation codons and prevent ribosome run-off during ultracentrifugation to separate 80S ribosome-mRNA complexes from untranslated mRNAs. Ribosome-bound mRNA was isolated and sequenced, and the relative abundance of each sequence was compared to its abundance in the input pool to determine a ribosome recruitment score (RRS) for each 5′-UTR (**Fig. 1c**). DART revealed up to 1,000-fold differences in ribosome recruitment that were highly reproducible (R^2^ = 0.94-0.95) among three independent replicates (**Fig. 1d)**. Thus, we established a robust method to determine the translation initiation activity of defined 5′-UTR sequences.

A caveat to the 80S/input calculation used to assess translation initiation activity is that sequences that destabilize 5′-UTRs in extracts could also lead to a reduction in 80S-bound RNA compared to the input pool. We therefore tested an alternative method of calculating ribosome recruitment scores by sequencing the untranslated mRNA from the top of the gradient in addition to the 80S fraction, RRS′ = 80S/(80S+mRNP) (**Supplementary Fig. 1a)**. These two metrics, RRS and RRS′, produced correlated but not identical results (R = 0.61 *p* < 2.2e-^16^, **Supplementary Fig. 1b**), suggesting that differential mRNA decay may affect the observed ribosome-bound pool to some extent. Because many effects on mRNA levels are likely to be downstream of changes in translation initiation^41–47^, the 80S/input calculation (RRS) is preferred. Importantly, each of the main conclusions reported below is consistent between calculation methods.

### Functional characterization of alternative 5’UTRs

Many eukaryotic genes express multiple mRNA isoforms that differ in their 5′-UTRs, which complicates the analysis of ribosome profiling data to identify translational control elements. Ribosome profiling measures ribosome occupancy per ORF^7^, but the same ORF frequently occurs in multiple transcripts with different regulatory regions^16–22^. Moreover, alternative 5′-UTRs can produce very large differences in translation activity as shown by reporter assays^23^ and by 5′-UTR variant specific polysome profiling^24–26^. Thus, the mRNA-specific translational efficiency inferred from ribosome profiling data may not accurately reflect the actual translation activity of any mRNA isoform (**Fig. 2a)**. We therefore used DART to directly compare the activity of expressed alternative mRNA isoforms whose 5′-UTRs differ by at least 10 nucleotides (**Supplementary Table 1**).

**Fig. 2.**
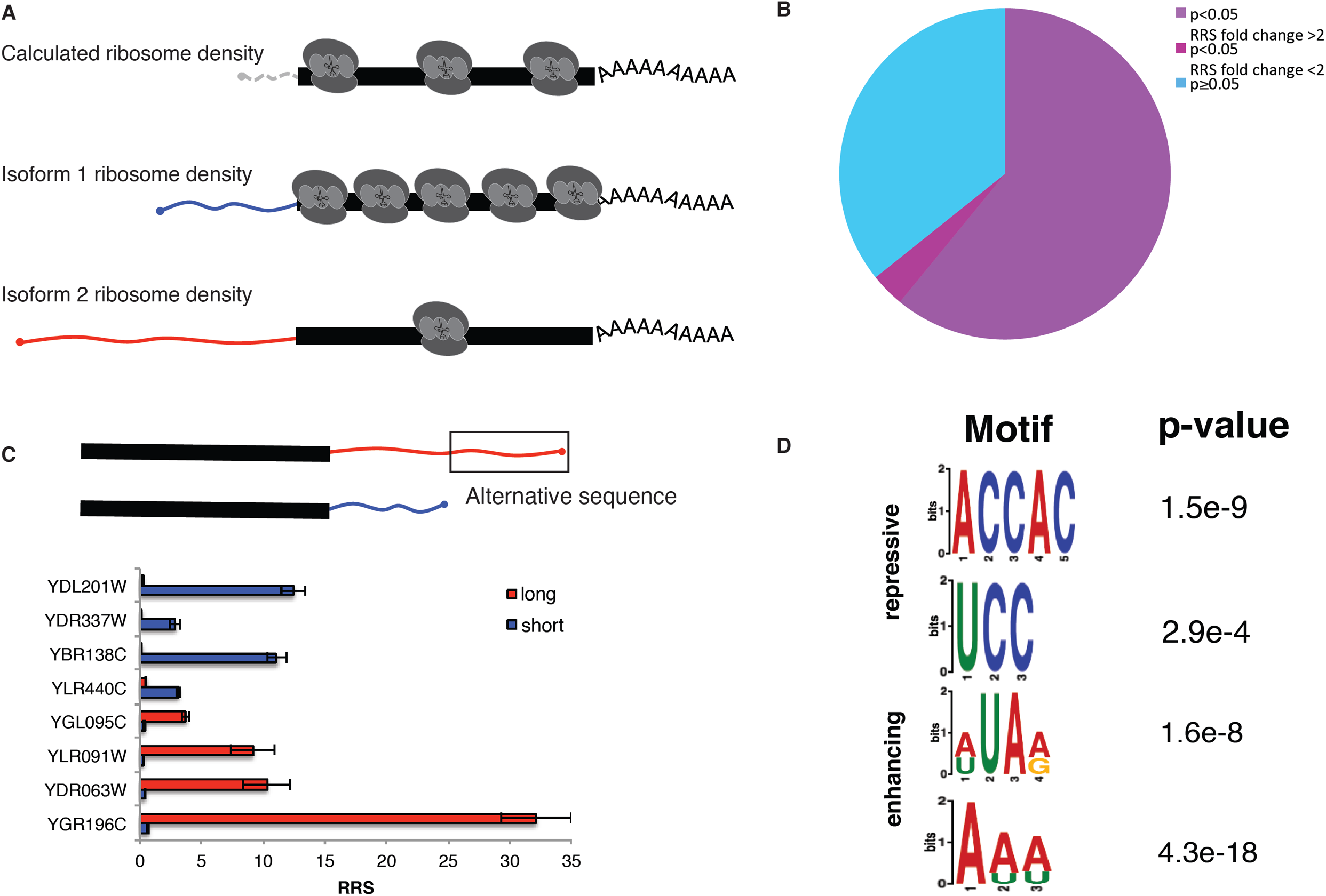
DART identifies translational silencers and enhancers within alternative 5′-UTRs. **(A)** 5′-UTR isoforms can confound the interpretation of ribosome density. **(B)** Most 5′-UTR isoforms tested significantly alter ribosome recruitment levels. Isoforms are separated by those that differ significantly (Bonferroni corrected p<0.05) and cause a >2-fold change in RRS (purple), those that differ significantly, but result in <2-fold change in RRS (magenta), and those that do not confer significant differences in RRS (blue). **(C)** Operationally defined enhancer and repressor regions. Additional sequence from longer isoforms that caused higher (enhancers) or lower (repressor) RRS levels were used as inputs to identify enriched motifs using DREME. **(D)** Top 2 enriched motifs from regions defined in 2C as identified by DREME. Repressive motifs (above) and enhancing motifs (below) are shown.

We tested 4354 alternative mRNA isoforms from 2064 genes and identified 1639 isoform pairs with reproducible differences in ribosome recruitment (*p* < 0.05, Bonferroni corrected two tailed t test) of which 843 differed by more than 3-fold (**Fig. 2b and Supplementary Table 2)**. These results are consistent with previous evidence that alternative 5′-UTRs differ in ribosome association in vivo^24–26^. Importantly, DART establishes the direct contribution of 5′-UTR sequences to differences in translation initiation and eliminates confounding effects of transcript localization and co-occurrence of alternative 3′-UTR sequences.

We leveraged these 5′-UTR isoform comparisons to identify novel translational enhancer and silencer elements, which were operationally defined as sequences present in a longer 5′-UTR variant that showed a higher RRS than a shorter alternative 5′-UTR of the same gene (enhancer) or lower RRS than the corresponding shorter alternative 5′-UTR (silencer). We identified 658 enhancer regions and 541 silencer regions by the criteria that their inclusion reproducibly changed RRS more than 2-fold (*p* < 0.001, n = 3 replicates) (**Supplementary Table 2)**. Of these, 72 enhancers and 67 silencers changed RRS by more than 10-fold (**Fig. 2c and Supplementary Fig. 2a**). Remarkably, sequence differences of only 10 nucleotides were sufficient to significantly alter translation initiation in many cases (**Supplementary Table 2**). To illuminate the mechanism of translational control, we looked for over-represented motifs within enhancers and silencers using DREME^27^ to identify short sequences that are over-represented. Intriguingly, the silencer regions were enriched in CCH motifs that resemble the C-rich motifs of the bottom 10% of all 5′-UTRs (**Fig. 2d and Supplementary Fig. 2c**), which suggests a common mechanism of translational repression. Overall, we found that 68% of genes with alternative 5′-UTRs showed significant differences in translation initiation between isoforms. These results demonstrate the broad potential for alternative promoter use, which generates alternative 5′-UTR isoforms, to regulate gene expression at the level of translation.

### Systematic testing of inhibitory 5′-UTR features

We exploited the throughput of the DART approach to systematically interrogate the effects of RNA secondary structure on translation initiation (**Fig. 3a**). Low-throughput assays have shown inhibitory effects of engineered stable stem-loop (SL) structures, which is partially consistent with global correlations between predicted 5′-UTR folding energy and translation activity^8,28^. However, there is conflicting evidence regarding the importance of SL position with respect to the 5′ cap or the AUG initiation codon (AUGi). We designed SL of varying stabilities (−5 to -31 kcal/mole) and inserted them into 20 natively unstructured 5′-UTRs (**Supplementary Fig. 3**). The complete set of 5′-UTR-SL constructs included 6 different stems, positioned every 3 nts across each 5′-UTR and extending 14 nts into the CDS, for a total of 1891 variants (**Supplementary Table 3**). All constructs were folded in silico to verify that a single structure was predicted for every sequence context.

**Fig. 3.**
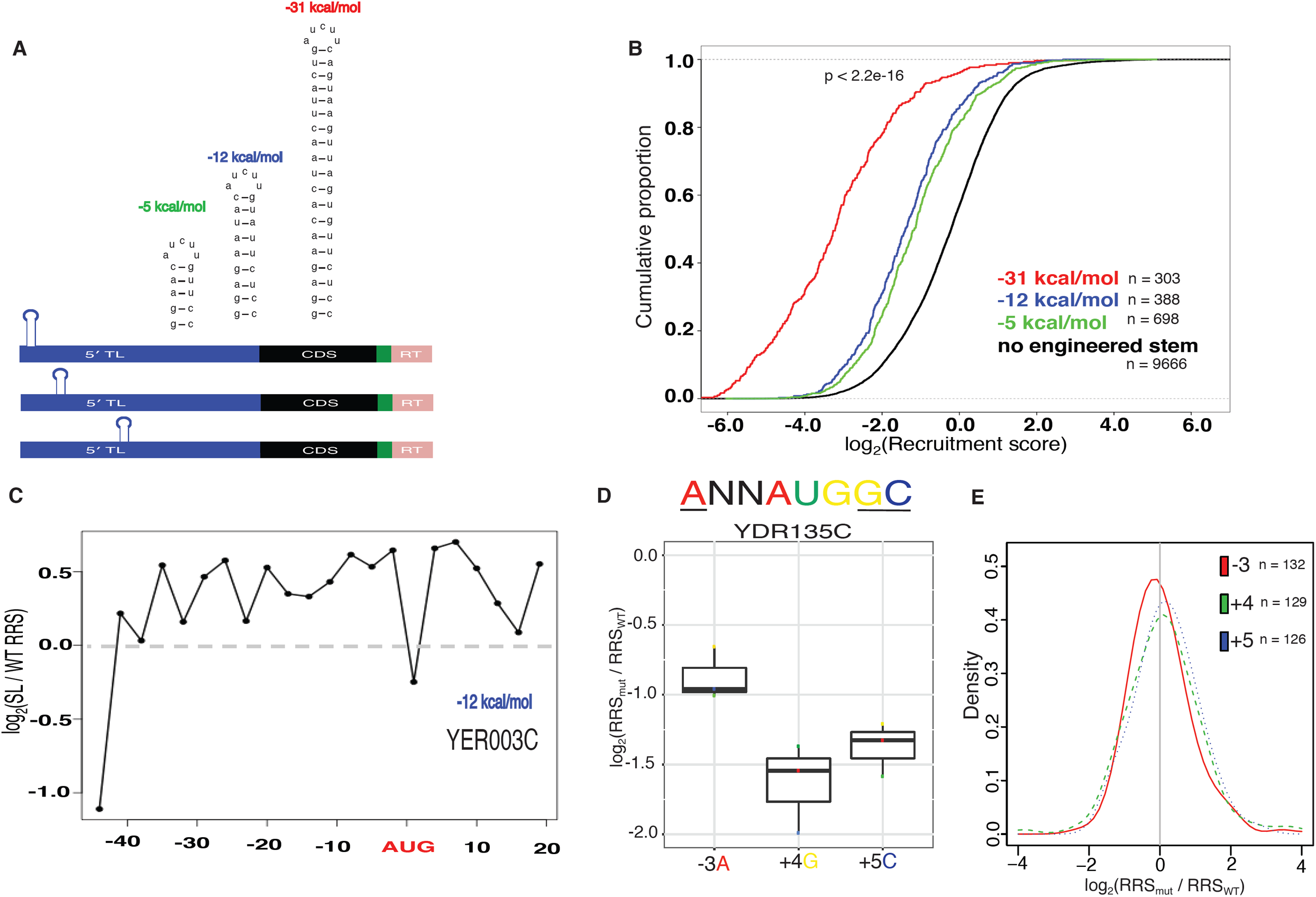
Inhibitory RNA features have highly context-dependent effects on initiation. **(A)** Design of stem loop constructs. Stem loops were designed to span a range of folding energies and were subsequently positioned every 3nts. An additional 3 stem loops were made by scrambling the stem sequences. **(B)** Engineered stem loops repress ribosome recruitment. All stems repress recruitment relative to sequences without an inserted stem (black) with the strongest stem (red) showing the greatest level of repression, followed by the intermediate stem (blue) and the weakest stem (green). **(C)** Individual genes show distinct effects of SL insertion. Relative recruitment for a single gene (YER003C) with the -12kcal/mol stem inserted. In this case, the stem is most repressive near the cap and at the start codon, but actually results in somewhat increased recruitment at other positions. **(D)** Genes show context dependent impacts of mutations to optimal AUGi context nucleotides. Relative recruitment score is shown for each mutation, with the identity of the mutated nucleotide indicated by the color of the point **(E)** Mutations at the -3 position are generally the most deleterious to ribosome recruitment. Distribution of relative RRS at each position.

The inserted SL structures inhibited translation initiation, with stronger inhibition observed with increasing stem stability (**Fig. 3b**). SLs with different sequences but the same folding energy inhibited to a similar extent as expected for a structure-based mechanism (**Supplementary Fig. 3**). Globally, the weaker SLs (−5 to -12 kcal/mole) inhibited ribosome recruitment most strongly at the cap-proximal position and actually enhanced translation initiation when positioned farthest from the cap (**Fig. 3c and Supplementary Fig. 3**). However, individual 5′-UTRs showed distinct positional effects of SL insertion (**Fig. 3c and Supplementary Fig. 3**), perhaps reflecting sequence context effects on RNA helicase activity^29^. The most stable SLs (−31 kcal/mole) strongly depressed translation, particularly from central positions (**Supplementary Fig. 2c**). However, as with most of the tested elements, we observe considerable variability in the effect of individual SL insertions **(Supplementary Figure 3d)**.

Current models of initiation invoke start codon context as an important feature in promoting translation. Consistent with this, uAUGs with optimal context are generally more inhibitory than those in a suboptimal context. However, AUGi context is not well correlated with ribosome occupancy making the extent of the role of start codon context unclear^30^. To test the role of the nucleotides surrounding AUGi directly, we designed constructs that altered the identity of one of three nucleotides that are either highly conserved or have previously been implicated in affecting translation^31^ (**Fig 3d**). We selected 50 5′-UTRs with the preferred ANN<u>AUG</u>GC context and mutated -3A, +4G and +4C to all other nucleotides to test 415 AUGi context variants (**Supplementary Table 4**). Overall, changes at the -3 position appear more deleterious than the other positions tested, although we do observe considerable variability (**Fig 3e**). However, while we observed many instances where mutation of preferred nucleotides reduced ribosome recruitment (**Fig 3d**), the effects appear to be highly context dependent and there are many examples where the changes actually improved recruitment (**Supplementary Fig. 3e**). Together these results with engineered SL structures and AUGi flanking sequences underscore the need to directly measure the contributions of mRNA features to translation activity within specific 5′-UTR contexts.

### A broad stimulatory role for initiation factor binding motifs in 5′-UTRs

5′-UTR elements that bind preferentially to eukaryotic initiation factors (eIFs) may function as translational enhancers. Consistent with this model, diverse viruses rely on high-affinity interactions between their 5′-UTRs and cellular initiation factors and/or ribosomes for efficient translation^32^. eIF4G, the scaffold subunit of the cap binding complex, is a prime candidate to mediate the activity of translational enhancer sequences. eIF4G contains three RNA binding domains that directly interact with mRNA and are essential for yeast growth^33,34^ although specific functional interactions between eIF4G and cellular mRNAs had not been characterized. We previously used RNA Bind-n-Seq (RBNS) to determine the RNA binding specificity of eIF4G1^35^. This competitive in vitro binding assay consists of mixing RNA libraries with different concentrations of protein, and sequencing the bound RNA^36^. Using a library of random 20mer RNA, we tested ∼87,380 distinct RNA 7mer motifs and showed that recombinant eIF4G1 preferentially binds to RNA sequences containing oligo-uridine (U). Consistent with this result, inserting oligo(U) into an unstructured RNA increased binding to eIF4G1 by 20-fold. Luciferase reporter assays supported a functional role for oligo(U) motifs in translation of five yeast mRNAs^35^.

Hundreds of yeast 5′-UTRs contain oligo(U) sequences, which are evolutionarily conserved among budding yeast species and enriched in genes with regulatory roles^35^. To directly test the impact of oligo(U) motifs on translation initiation, we performed DART on native 5′-UTR sequences with or without oligo(U) motifs. We synthesized a pool of capped mRNAs consisting of all yeast 5′-UTR sequences ≤94 nt long that contain U≥7 (168 in total, **Supplementary Table 5**) together with their start codons and some coding sequence. The pool included matched controls for each 5′-UTR in which the oligo(U) motif was replaced with a CA repeat of equal length (**Fig. 4a**). In all 3 replicates, oligo(U)-containing mRNAs were recruited more efficiently to the ribosomal fraction than their matched mutant counterparts (**Fig. 4b**). Of the 13 5′-UTR sequences whose ribosome recruitment differed significantly (p < 0.05 by 2-tailed paired t-test) between the wild type and mutant variants, none were recruited better with the oligo(U) mutated to oligo(CA) (**Fig. 4c**). Five of these 5’UTRs were cloned upstream of nanoluciferase and their translation measured in yeast lysates. Only one of the five 5′-UTRs (YMR158W, 56nt 5′-UTR) increased luciferase production when mutated to oligo(CA) (**Fig. 4d**). This 5′-UTR sequence is predicted to form a stem-loop that sequesters the oligo(U) sequence (**Supplementary Fig. 4**), which would likely preclude eIF4G binding^35^, and could have additional inhibitory effects (**Fig. 3**). The CA mutation disrupts this structure, which could potentially explain the enhanced luciferase production from the mutant 5′-UTR. These findings establish a broad stimulatory role for eIF4G binding motifs in 5′-UTRs and illustrate the potential for DART to illuminate the function of putative RNA regulatory elements identified by other high-throughput approaches.

**Fig. 4.**
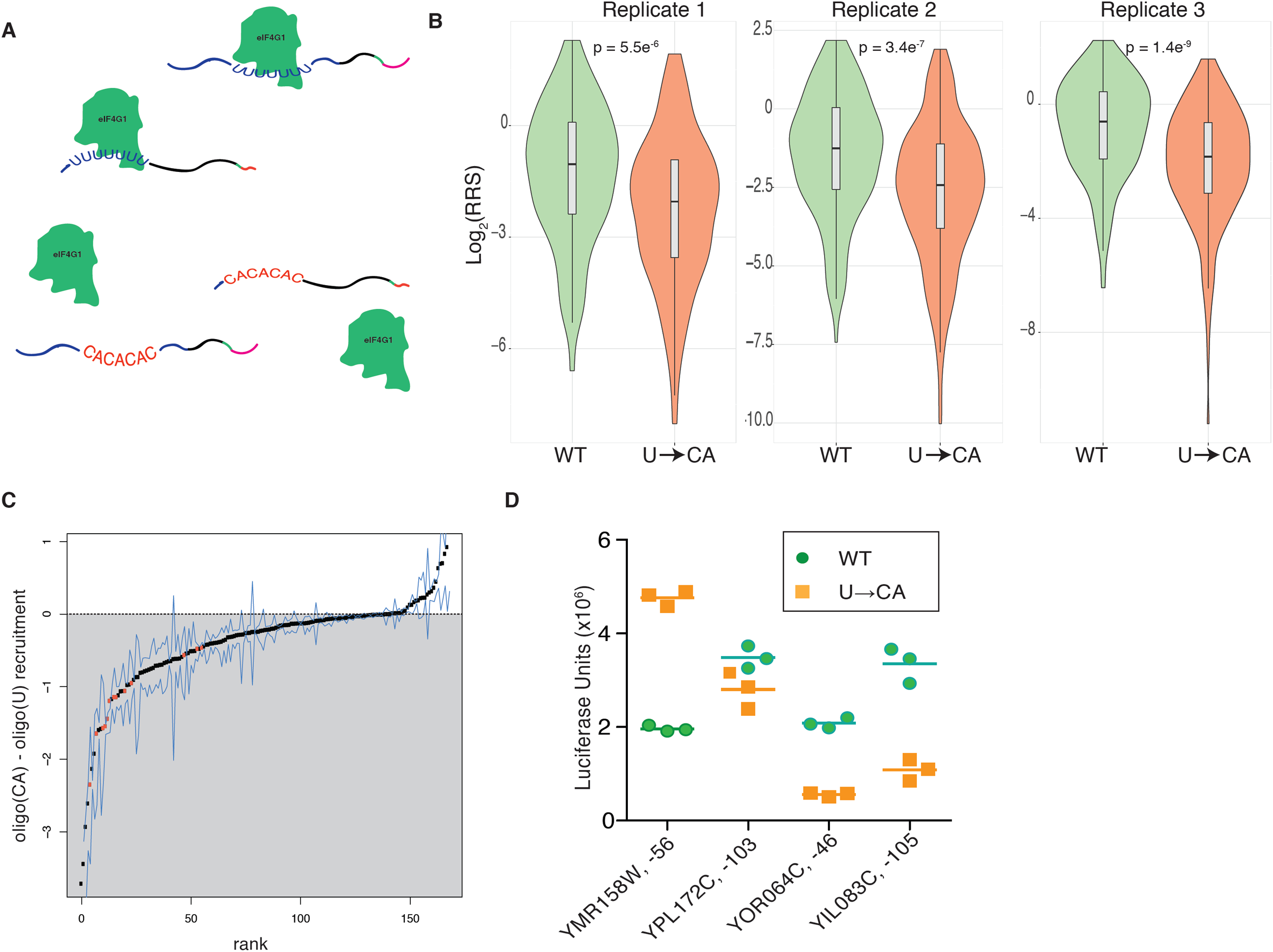
eIF4G binding motifs promote translation initiation. **(A)** Schematic of eIF4G1 binding to designed RNA pool including both WT oligo(U) motifs (blue) and U->CA mutants (red). **(B)** Violin plots of ribosome recruitment score distributions for 5′-UTRs containing U≥7, and the matched controls, for 3 independent replicates. **(C)** Ranked changes in ribosome recruitment scores between oligo(U) 5′-UTRs and matched U->CA mutants. Each dot is the mean of all replicates (excluding values that did not meet read coverage thresholds). Blue lines indicate the minimum and maximum value across the replicates. 5′-UTRs with a statistically significant (Bonferroni-corrected two-tailed t-test p < 0.05) difference between WT and mutant variants are colored orange. **(D)** NanoLuc activity from *in vitro* translation extracts programmed with capped and polyadenylated NanoLuc mRNA bearing the indicated 5′-UTR. Each point represents an independent experiment, and is the mean of 3 technical replicates. Horizontal lines indicate the mean of the three independent replicates. For each 5′-UTR, the difference between the means of the WT and mutant NanoLuc activity is statistically significant with p<0.05 (two-tailed student’s t-test).

## Discussion

We have developed direct analysis of ribosome targeting (DART) for high-throughput functional testing of 5′-UTR variants to illuminate the mechanisms underlying widespread translational control. The low density of ribosomes observed even on highly translated mRNAs suggests that translation initiation, as opposed to elongation or termination, is rate-limiting for most genes^37^. However, local differences in elongation speed clearly affect the abundance of ribosome-protected footprints thereby confounding interpretation of ribosome profiling data^38^. DART enables greater mechanistic insight by effectively isolating translation initiation from other processes and 5′-UTR effects from other co-varying mRNA regulatory elements.

Quantitative analysis of translation initiation in vitro has begun to reveal distinct initiation mechanisms used by different mRNAs^29,39^. The characterization of translation inhibition by RNA structures presented here illustrates the importance of testing many mRNA sequences. The broad, but weak, correlations previously observed between predicted 5′-UTR folding energy and translation activity^8,9^ can now be understood as the probable effect of highly context-dependent effects of RNA structures. Sequence context effects may similarly account for the failure of RNA folding to predict more than 36% of the observed variance in translational repression following inactivation of the mRNA helicase, Ded1^40^. It will be interesting to extend previous in vitro characterization of Ded1-dependent mRNAs to a much broader sample of 5′-UTRs^29^. The capacity of DART to quantify translation initiation on thousands of defined 5′-UTR variants in biochemically accessible cell lysates will be broadly useful to elucidate fundamental translational mechanisms.

We note that 5′-UTR-dependent changes in translation activity in vivo, as determined by measuring steady-state reporter protein activity normalized by steady-state mRNA levels, are consistently smaller than the 5′-UTR-dependent fold changes in protein production observed pre-steady state in vitro^23^. This is the expected result if poorly initiated mRNAs are more rapidly degraded, as suggested by destabilization of reporter mRNAs with defects in translation initiation^41–43^ and by global anti-correlations between TE and mRNA half-life^44–47^. Combining DART with direct assessment of mRNA half-lives in intact as well as ribosome-depleted lysates will enable a complete characterization of the interplay between translation initiation and mRNA decay.

Here we used DART to characterize the translational activity of alternative 5’UTR isoforms, revealing functional differences between 1639 isoform pairs (**Fig. 2b and Supplementary Table 2**). This is likely to be a broadly used regulatory strategy as many eukaryotic genes express multiple mRNA isoforms due to alternative transcription start site (TSS) selection, which is highly regulated^16–20^. Notably, yeast transcript isoform analysis across different environmental conditions identified more than 1200 genes with regulated alternative 5′-UTRs^21^ and alternative TSS selection contributes more to mammalian mRNA isoform diversity than alternative splicing in some tissues^22^. Tumor-specific changes in promoter usage have also been observed in dozens of different cancer types^48^. DART will be broadly useful to determine how sequence differences between alternative 5′-UTRs affect protein production, which is critical for predicting the functional output of normal or pathological transcriptional regulation that changes the type of mRNA produced as well as the number of mRNA molecules.

By strategic design and high-throughput testing of 5′-UTR mutants, DART makes it possible to pinpoint the functional nucleotides within candidate regulatory elements implicated as translational enhancers or silencers. We used DART to systematically probe the effect of oligo uridine motifs previously identified as preferred binding sites for the translation initiation factor eIF4G1^35^. Our results established a general stimulatory role for this 5′-UTR element and identified interesting outliers that may reflect RNA context effects that can be probed in subsequent DART experiments. A related application of DART will allow rapid functional screening of 5′-UTR SNPs that have been linked to diverse disease phenotypes in genome-wide association studies (GWAS). Current efforts to identify the causal variants, which are potential targets for new therapies, are largely restricted to non-synonymous changes within protein coding sequences. The DART method will enable systematic identification of 5′-UTR variants that are likely to be damaging by dysregulating protein levels. Finally, DART will be useful to optimize 5′-UTRs for protein production in a variety of settings, including in therapeutic mRNAs.

## Methods

### 5′-UTR Pool Design and Synthesis

Each sequence in the pool consisted of ≤122 nucleotides of 5′-UTR sequence followed by at least 24 nucleotides of coding sequence followed by a randomized 10 nucleotide unique identifier barcode and an adaptor sequence used for priming reverse transcription and Illumina sequencing (**Fig 1a**). For 5′-UTR annotations, 5′-UTR abundances were calculated from^49^. Each 5′-UTR was merged with it’s most abundant neighbor within 10 nts in either direction such that no two 5′-UTRs were within 10 nts of each other. We then imposed the following requirements (1) 5′-UTRs must be expressed within 25% of the mode abundance for a given 5′-UTR, and (2) 5′-UTRs must make up at least 5% of the total abundance for that ORF, unless the mode was <5% of the total, in which case we used the mode. Upstream AUGs within 5′-UTR sequences were mutated to AGT such that the first AUG encountered by a scanning pre-initiation complex moving 5′ to 3′ would be the annotated AUGi. Designed oligos were purchased as a pool (Twist Bioscience) and PCR amplified with RN7 and RN8 (**Supplementary Table 6**). RNAs were produced by runoff T7 transcription from gel-purified DNA template. RNAs were 5′ capped using the Vaccinia Capping System (NEB M2080S) and 3′ biotinylated using the Pierce RNA 3′end biotinylation kit (Thermo Scientific 20160).

### In Vitro Translation

Yeast translation extracts were made as described in (Rojas-Duran and Gilbert, 2012) from YWG1245 (MATa trp1Δ leu2-3,112 ura3-52 gcn2Δ ::hisG PGAL1 -myc-UBR1 ::TRP1 ::ubr1, pRS316 <URA3>) cultured in liquid YPAD (1% yeast extract, 2% peptone, 2% glucose, 0.01% adenine hemisulfate). Approximately 40 pmoles of mRNA were added per 2 mL translation reaction. Reactions were incubated with 0.5mg/mL cycloheximide at 26**°**C for 30 minutes in a shaking thermomixer.

### 80S isolation

Translation reactions were loaded onto 10%-50% sucrose gradients in polysome lysis buffer 20 mM HEPES-KOH pH 7.4, 2 mM MgAc, 0.1 M KAc, 0.1 mg/mL cycloheximide, 1% TritonX-100 followed by centrifugation at 27,000 rpm in a Beckman SW28 rotor for 3 hours. Gradients were fractionated from the top down using a Biocomp Gradient Station (Biocomp Instruments) with continual monitoring of absorbance at 254 nm. Fractions corresponding to the 80S peak were pooled and RNA extracted using phenol/chloroform followed by isopropanol precipitation.

### Library preparation

Biotinylated RNA was resuspended in binding buffer (0.5M NaCl, 20mM Tris-HCl pH 7.5, 1mM EDTA) and isolated using Hydrophilic Streptavidin Beads (NEB S1421S). Isolated RNA was reverse transcribed using the barcoded primer OWG921 and Superscript III (Invitrogen 18080093). Gel-purified cDNA products were ligated to the adapter oWG920 (**Supplementary Table 6**) using T4 RNA ligase 1 (NEB M0437M). cDNA cleanup was performed using 10 µl MyOne Silane beads (Thermo Scientific 37002D) per sample. Libraries were then PCR amplified with primers RP1 and OBC (**Supplementary Table 6**) and sequenced on a HiSeq 2500.

### Read processing

Reads were first demultiplexed by their library-level barcodes. PCR duplicates were collapsed using FastUniq. Output sequences were trimmed of the 5’ decamer barcode FastX trimmer. Reads were uniquely mapped to the sequences within the synthetic pool using STAR with the command line options < --outSAMstrandField intronMotif --alignIntronMin 200 > Any reads that failed to align were excluded from further analysis. FPKMs were quantified using cufflinks.

## Supplementary Figure Legends

**S1.**
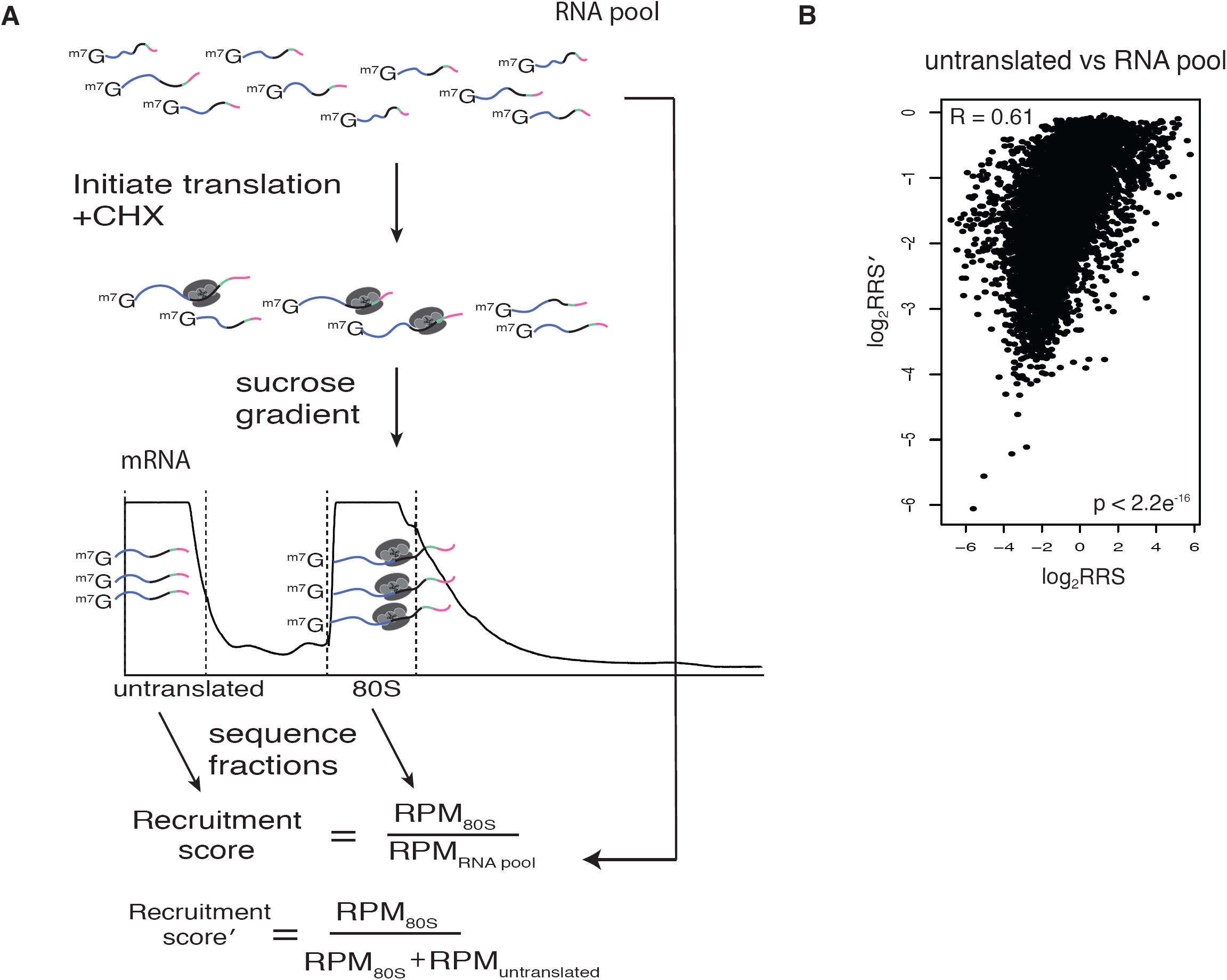
Alternative RRS calculations. **(A**) DART workflow with all sequenced fractions. The RNA pool, input, and untranslated fractions can all be sequenced to give different RRS values that include RNA stability effects. **(B) Correlation between RRS calculation methods**.

**S2.**
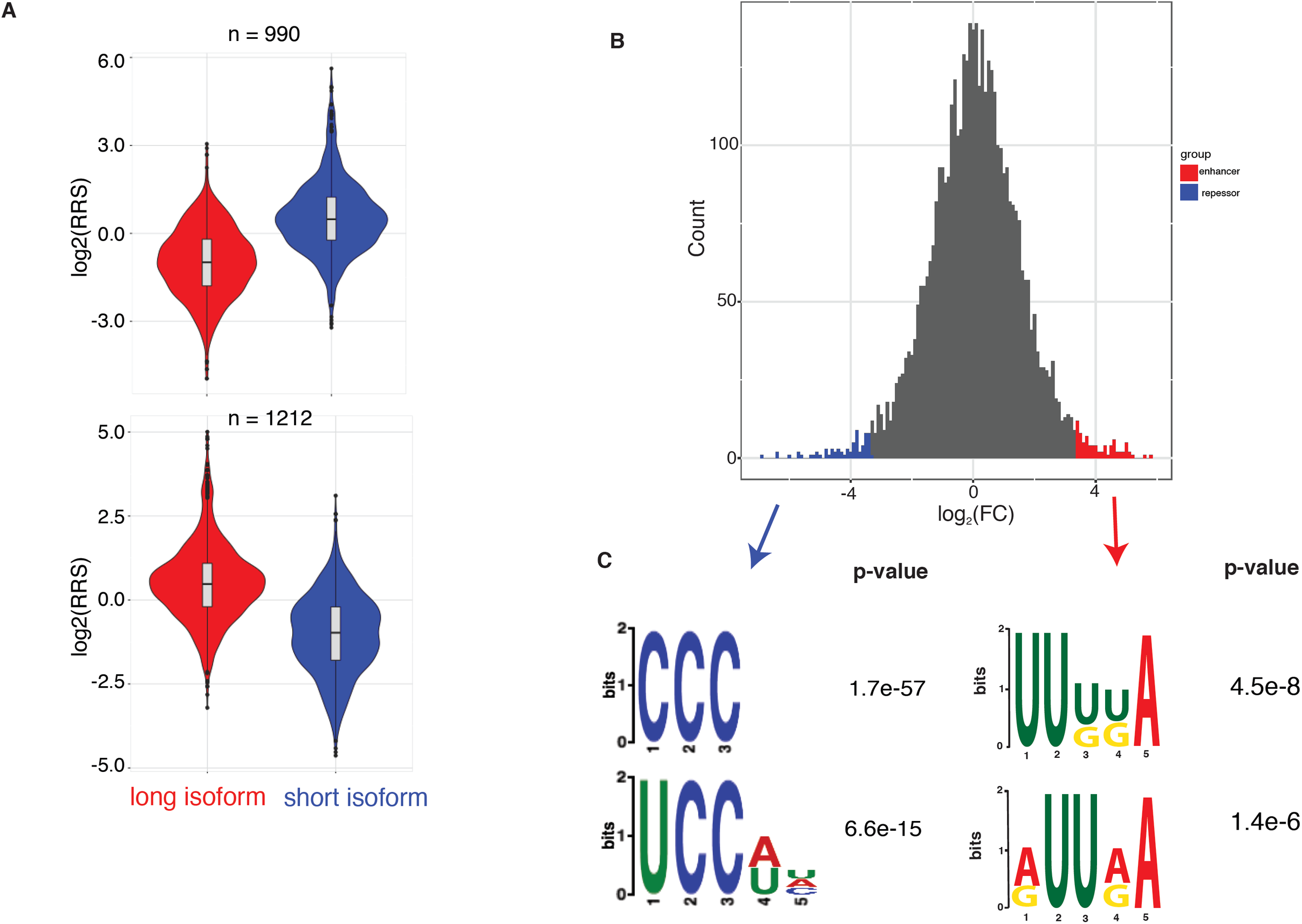
Translational enhancers and repressors identified by DART. **(A)** 5′-UTR isoforms used for motif analysis. Alternate sequence from longer 5′-UTR isoform that show lower or higher ribosome recruitment were used to search for enriched motifs against the background of all differential sequences. **(B)** Top and bottom decile of RRS used to search for enriched motifs. **(C)** Identified repressor (left) and enhancer (right) elements. Motifs identified from top and bottom decile of RRS shown in S1B.

**S3.**
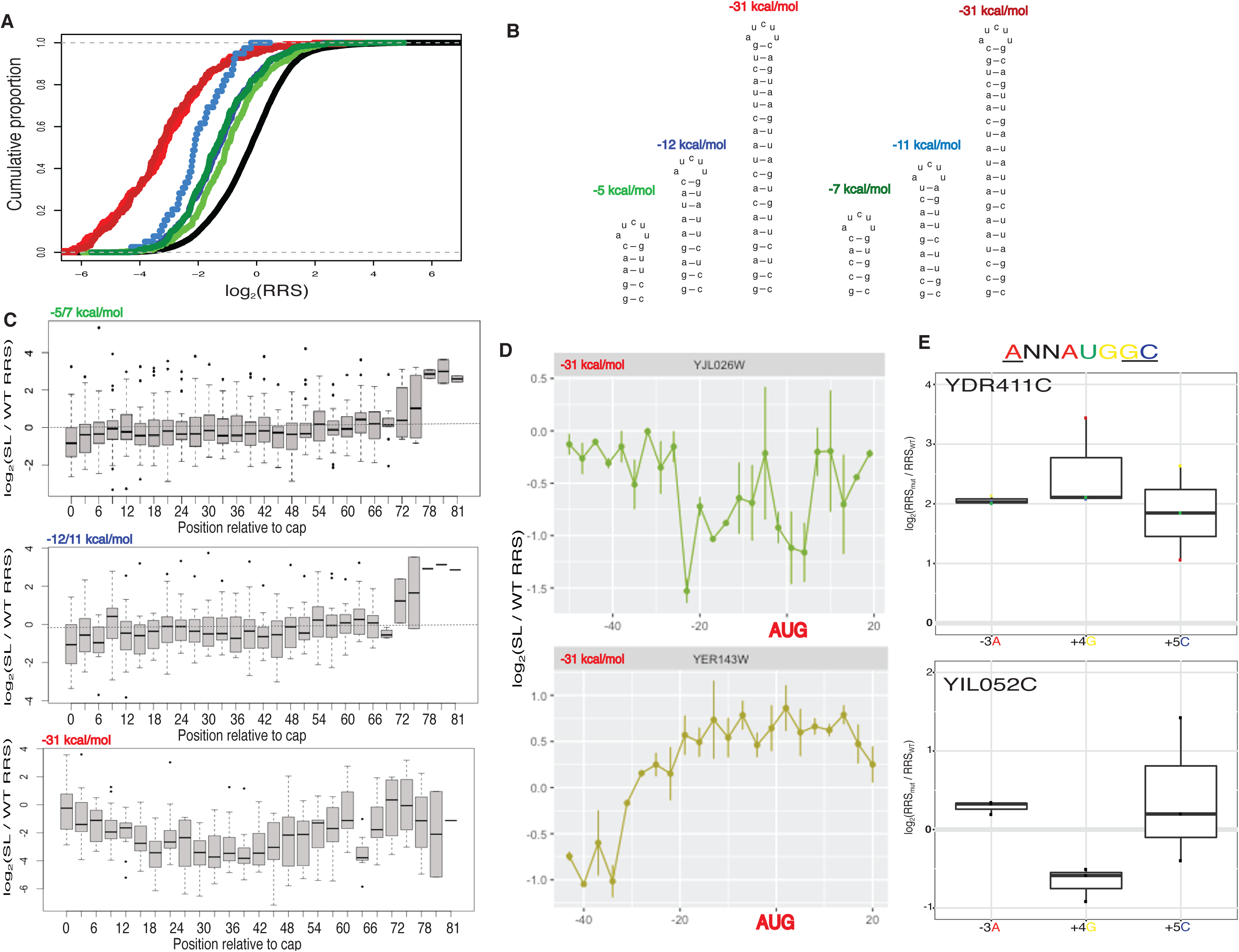
RNAs show a range of responses to inhibitory elements. **(A)** Designed stem loop constructs. Designed stems with sequence-scrambled counterparts. Color scheme the same as in (B). **(B)** Stem loops with equivalent strength repress ribosome recruitment similarly. Designed stems loops and their scrambled sequence counterparts repress recruitment to a similar degree. **(C)** Designed stems exhibit some position-dependent effects. Box plot of ribosome recruitment levels for the -5 kcal/mol stem relative to the parent constructs with no SL. Ribosome recruitment with inserted stems of a given strength plotted by position relative to the mRNA cap. Stems inserted farthest from the cap generally promote ribosome recruitment. **(D)** Stem loops exhibit context-dependent effects on RRS. Relative ribosome recruitment by position for two individual gene contexts (YJL026W and YER143W). **(E)** Start codon context mutations. Genes YDR411C and YIL052C respond differently to start codon context mutations.

**S4.**
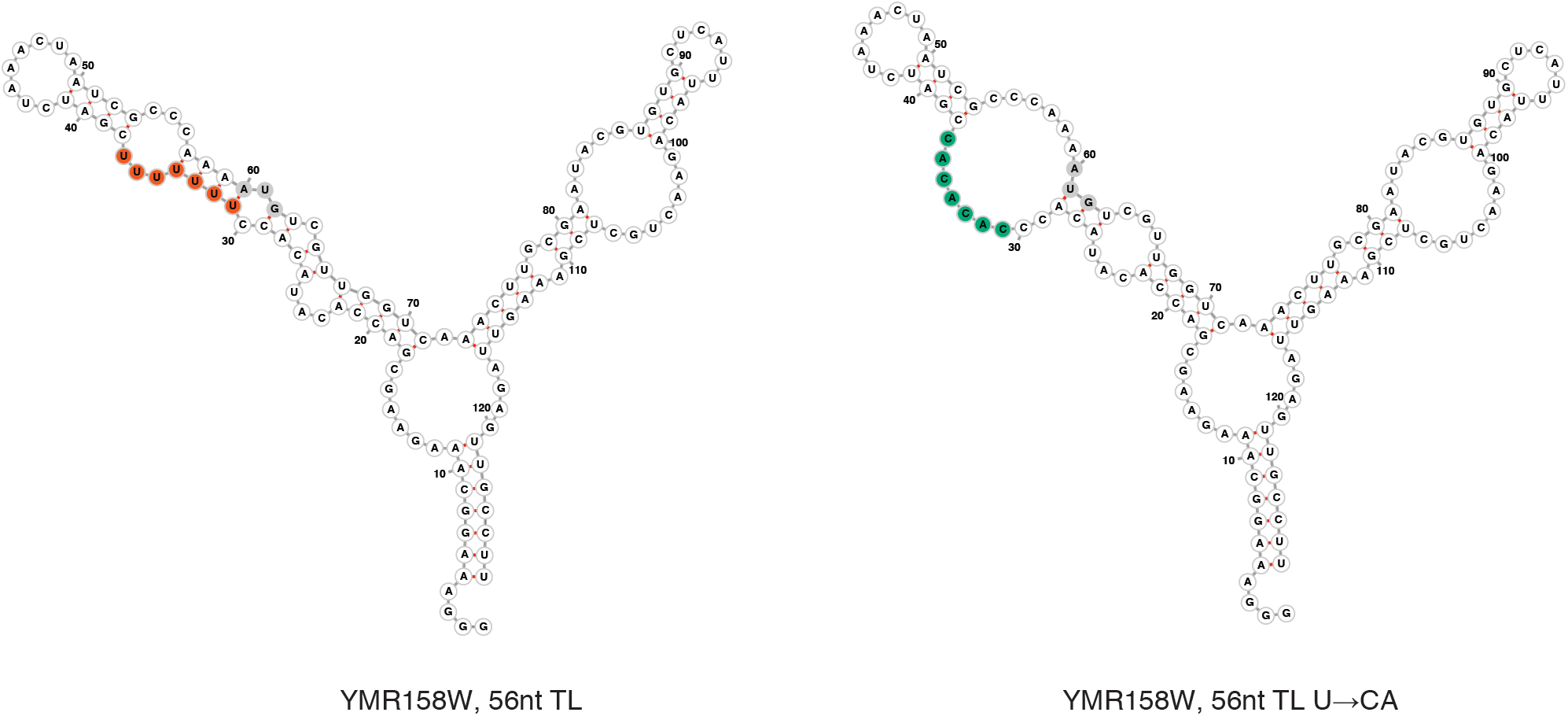
Mutation in the oligo(U) motif in YMR158W disrupt base pairing. Folding predictions in WT and oligo(CA) mutated YMR158W.

**S5.**
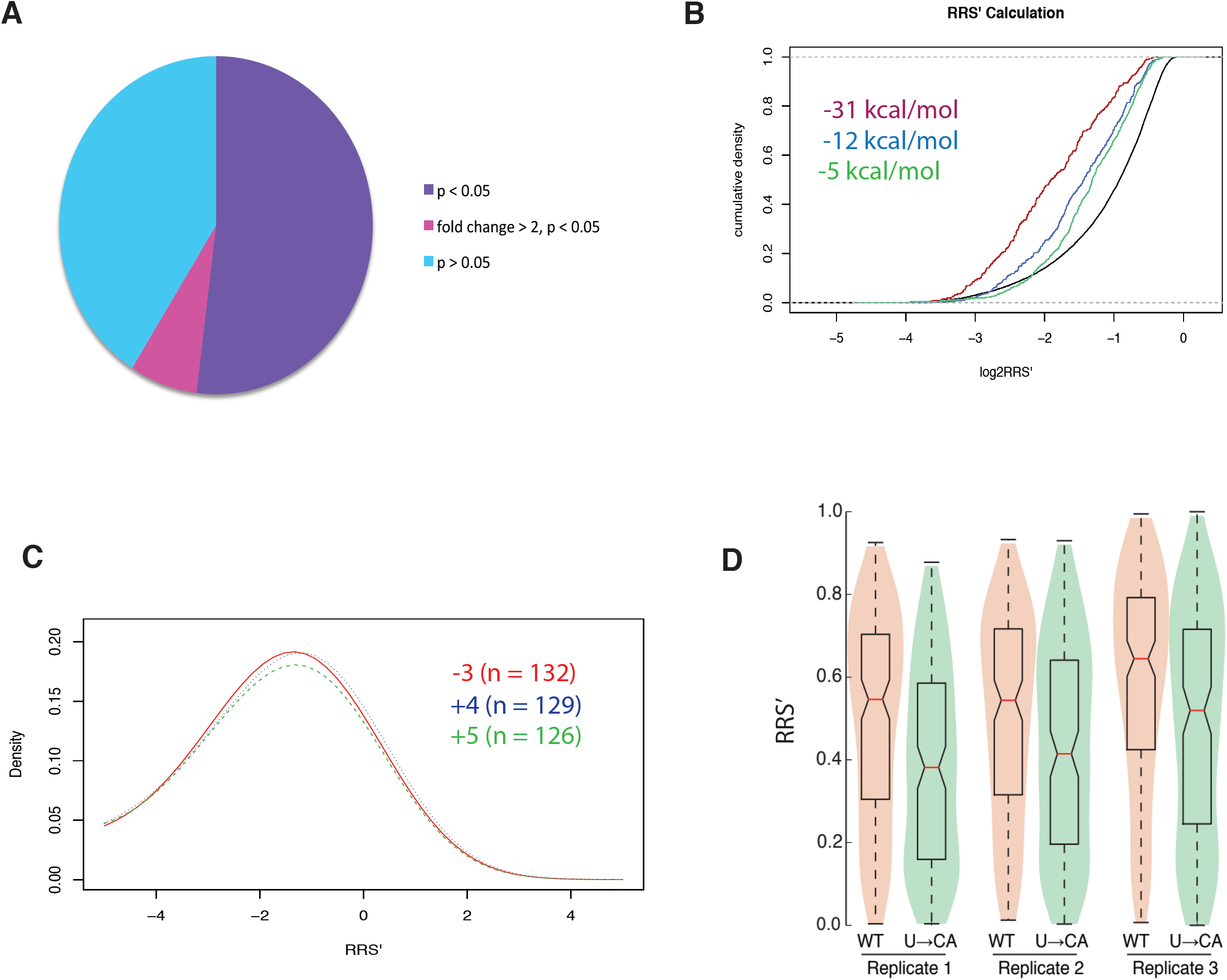
Features tested using alternative RRS calculations. **(A)** Most 5′-UTR isoforms tested significantly alter ribosome recruitment levels using the RRS′ calculation. **(B)** Stronger stems show lower RRS′ values. **(C)** Mutating conserved AUGi context nucleotides reduces RRS′, although the effects are highly variable. **(D)** Mutating oligo(U) sequences results in lower RRS′.

**S6.**
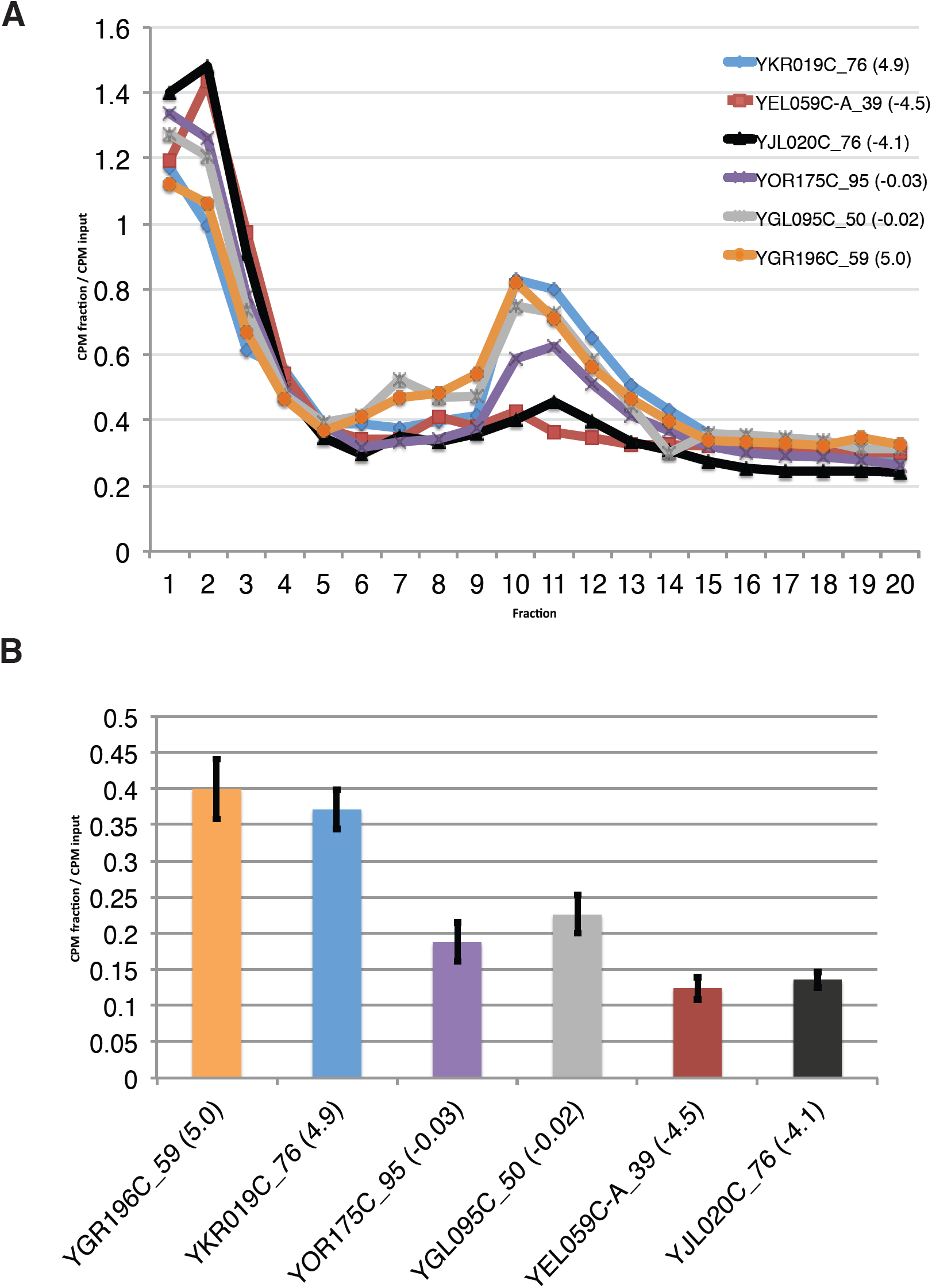
Low-throughput validation of DART. 5′-UTR sequences spanning a range of RRS values (indicated in parenthesis) were radiolabeled and incubated in yeast translation extract in the presence of cycloheximide. Reactions were loaded onto a sucrose gradient. **(A)** Counts in each fraction are reported relative to counts from the input RNA. **(B)** Quantification of three independent replicates shown.

